# Frontal cortex function derives from hierarchical predictive coding

**DOI:** 10.1101/076505

**Authors:** William H Alexander, Joshua W Brown

## Abstract

The frontal lobes are essential for human volition and goal-directed behavior, yet their function remains unclear. While various models have highlighted working memory, reinforcement learning, and cognitive control as key functions, a single framework for interpreting the range of effects observed in prefrontal cortex has yet to emerge. Here we show that a simple computational motif based on predictive coding can be stacked hierarchically to learn and perform arbitrarily complex goal-directed behavior. The resulting Hierarchical Error Representation (HER) model simulates a wide array of findings from fMRI, ERP, single-units, and neuropsychological studies of both lateral and medial prefrontal cortex. Additionally, the model compares favorably with current machine learning approaches, learning more rapidly and with comparable performance, while self-organizing representations into efficient hierarchical groups and managing working memory storage. By reconceptualizing lateral prefrontal activity as anticipating prediction errors, the HER model provides a novel unifying account of prefrontal cortex function with broad implications both for understanding the frontal cortex and building more powerful machine learning applications.

## Introduction

The frontal lobes are central to volition and higher cognitive function, especially goal-directed behavior^1–3^. Recent work has highlighted reinforcement learning, ^4–6^ performance monitoring ^7,8^, and hierarchical abstraction and working memory ^9–11^ as key elements of frontal function, often under the framework of cognitive control^12^. Considering the range of methods and perspectives applied to investigating prefrontal cortex (PFC), there is a clear need for a common framework for interpreting the variety of functions assigned to the frontal lobes.

Within the past decade, predictive coding has emerged as just such a potentially unifying framework for understanding the organization and function of the brain ^13^. Hierarchical predictive coding, as well as related approaches including free energy ^14^ and Hierarchical Bayesian Inference ^15^, generally treat bottom-up processing of information in the brain as a source of evidence that must be “explained away” by top-down processes carrying information regarding the likely causes of sensory information. In the predictive coding framework, top-down processes provide predictions from superior hierarchical levels to inferior levels, while residual prediction errors, i.e., input that cannot be accounted for by the predictions supplied by top-down processes, are carried from inferior levels to superior levels. This motif of top-down predictions and bottom-up prediction errors repeats through successive hierarchical iterations, forming a sophisticated processing stream composed of “dumb processes that correct… error in the multi-layered prediction of input.”^13^. Predictive coding accounts have achieved great success in accounting for effects related to the processing of sensory input ^16–22^. Given this success in accounting for the structure and function of the brain in early sensory areas, it has been suggested^13^ that the predictive coding framework might be extended to account for the organization of brain regions underlying sophisticated cognitive processes, especially the frontal lobes. To date, however, this proposed extension has remained largely hypothetical, and it remains an open question as to whether frontal lobe function can be accommodated within the predictive coding framework.

There are several reasons to believe that predictive coding formulations may indeed map well to PFC in addition to primary sensory areas. PFC is generally considered to be organized hierarchically along a rostrocaudal abstraction gradient^9,10,23,24^, with rostral regions coding for abstract rules and task sets, while caudal regions represent concrete stimulus-response associations. Significant portions of PFC are specialized for reporting error as a deviation from predicted events ^7,25^, and distinct regions within medial PFC (mPFC) appear to encode error at different levels of abstraction^26,27^, while regions within dorsolateral PFC (dlPFC) appear to encode hierarchical task set information ^23^ and to contextualize behavioral responses based on a learned model of the environment^10,24^.

In recent work^28^ we proposed the *error representation hypothesis,* formalized in the Hierarchical Error Representation (HER) computational model, of the functions and interactions of mPFC/dlPFC. We began with the PRO model of mPFC, which simulates a wide range of empirical findings as resulting from prediction errors calculated within mPFC ^7,25^. We then proposed that the mPFC prediction errors further train dlPFC to represent items in working memory that reliably predict the prediction errors generated by mPFC. These “error prediction” signals in turn are deployed to reduce subsequent prediction errors. Residual errors - those that cannot be fully predicted at a given level - act as a “proxy” outcome for higher levels of a mPFC/dlPFC hierarchy, and these proxy outcomes may in turn be the targets for further prediction and error computations. The result is a self-organizing hierarchical network that learns, maintains, and flexibly switches working memory representations as a product of learning to minimize prediction error. (Figure 1; supplementary material/methods). Notably, the HER model reconceptualizes working memory function as a product of learning to maintain representations of predicted errors, which in turn minimize prediction errors. In the framework of predictive coding, items in working memory in the HER model reflect hypothetical causes of observations that are selected based on bottom-up error signals, and the predictions of which are refined in order to explain away error at lower hierarchical levels. Hierarchical representation in the model therefore emerges as each hierarchical level identifies the most likely causes of residual errors reported by lower levels, and the degree to which a given hierarchical level influences the processing of a lower level is proportional to the amount of additional error that is accounted for given a hypothetical cause (supplementary materials/methods).

**Figure 1.**
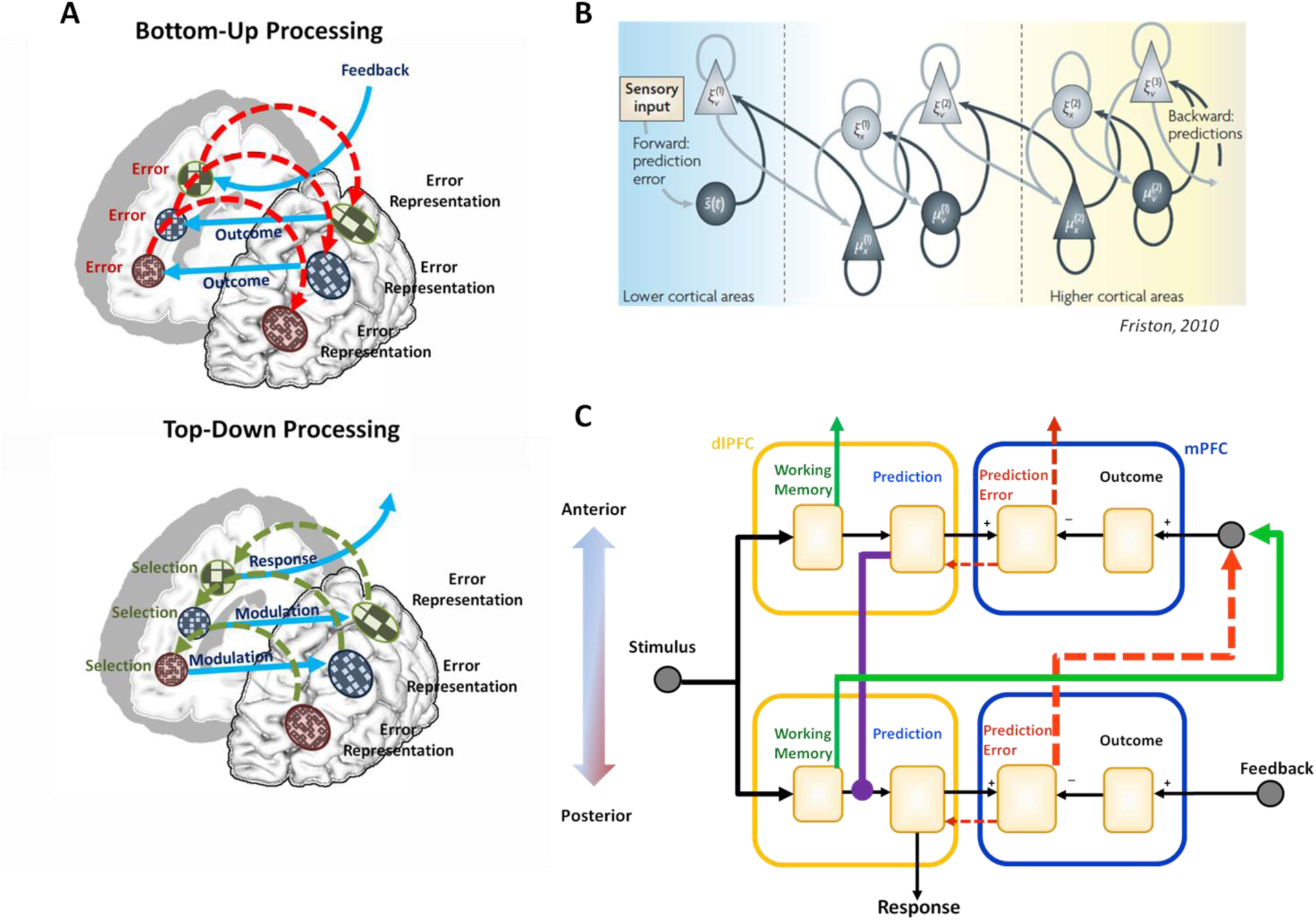
Predictive Coding in Prefrontal Cortex. A) In the HER model, information is passed to hierarchical levels through bottom-up and top-down pathways. In the bottom-up paths (top), regions in mPFC compute an error signal as the discrepancy between the expected and actual output of inferior hierarchical levels. Error signals generated by mPFC train error predictions in lateral PFC which are associated with task stimuli that reliably precede them. Following training, learned representations of error predictions are elicited by task stimuli and actively maintained in dlPFC for as long as they have predictive value. In the top-down pathway (bottom), error predictions are passed from superior hierarchical levels in order to successively modulate predictions made at inferior levels. B) The organization of the HER model is similar to formulations of predictive coding and free energy previously used to explain results from early sensory processing areas and hypothesized to extend into the frontal lobes. C) A detailed circuit diagram of the HER model shows bottom-up (red and green) and top-down (violet) pathways, as well as the working memory gating mechanism that allows information to be maintained over extended durations. The connections match known neuroanatomy^**29,30**^.

Computational simulations of the HER model have demonstrated its ability to learn complex cognitive tasks in a manner comparable to human performance, both in terms of behavioral markers of learning as well as the speed at which such tasks were acquired ^28^. Furthermore, the HER model performs well compared to state-of-the-art “deep learning” models such as LSTM ^28^ (cf. Table 1). The model’s ability to perform these tasks is noteworthy considering that it is composed of a repeated motif of relatively “dumb processes” organized hierarchically: individual hierarchical levels instantiate simple RL learners that receive feedback in the form of error signals generated by lower levels, and whose predictions serve to modulate lower level predictions.

**Table 1:** The HER model performs comparably to several out-of-the-box machine learning algorithms using the data sets described by Meyer et al., 2003^37^. Dark grey cells reflect those tasks in which the HER model equaled or exceeded the best classifier tested in that paper for various definitions of error (see Online Methods). In particular, the model tends to perform well on data that are principally categorical in nature, while performing less well on metric data sets. This is consistent with the design emphasis of the HER model on learning rules and task-sets required for performing typical cognitive tasks.

|  |  | Meyer et al., (2003) |  | HER MODEL |  |  |  |
| --- | --- | --- | --- | --- | --- | --- | --- |
|  |  | Pct. Error<br>(MEDIAN) | Pct. Error<br>(MEAN) | Pct. Error 1<br>(MEDIAN) | Pct. Error 2<br>(MEDIAN) | Pct. Error 1<br>(MEAN) | Pct. Error 2<br>(MEAN) |
| Categorical Data | Breast Cancer | 2.92% | 3.14% | 4.29% | 4.29% | 5.07% | 4.39% |
|  | HouseVotes | 0% | 0% | 4.55% | 2.33% | 4.23% | 2.78% |
|  | Heart1 | 13.33% | 13.67% | 16.67% | 12.70% | 18.19% | 12.35% |
|  | cards | 10.94% | 11.44% | 14.49% | 10.14% | 14.66% | 11.02% |
|  | hepatitis | 0% | 0% | 17.65% | 12.13% | 18.29% | 11.45% |
|  | tictactoe | 0% | 0.14% | 0.00% | 0.00% | 0.30% | 0.20% |
|  | chess | 0.31% | 0.49% | 1.25% | 0.31% | 1.97% | 0.61% |
|  | monk3 | 1.75% | 1.07% | 1.75% | 1.75% | 1.19% | 1.07% |
|  | promotergene | 9.09% | 10.96% | 9.09% | 8.33% | 10.69% | 8.11% |
|  | titanic | 20.81% | 21.09% | 20.81% | 20.36% | 21.17% | 20.29% |
| Metric Data | Sonar | 15% | 12.69% | 19.05% | 14.29% | 20.70% | 15.14% |
|  | PIDiabetes | 22.08% | 22.60% | 23.68% | 20.78% | 24.84% | 21.42% |
|  | liver | 27.02% | 29.11% | 31.43% | 27.78% | 32.30% | 28.65% |
|  | musk | 6.12% | 5.98% | 24.49% | 16.67% | 25.19% | 17.35% |
|  | credit | 22.77% | 23.12% | 25.25% | 20.90% | 25.88% | 20.60% |
| Artificial Data | Circle | 2.50% | 2.66% | 20.80% | 20.50% | 21.53% | 20.82% |
|  | Spirals | 0.50% | 0.17% | 44.35% | 42.60% | 44.00% | 42.39% |
|  | twonorm | 2.70% | 2.82% | 4.50% | 3.80% | 5.11% | 4.03% |
|  | threenorm | 13.70% | 15.76% | 19.40% | 17.30% | 20.08% | 17.05% |
|  | ringnorm | 2.90% | 3.58% | 27.70% | 24.00% | 28.83% | 24.02% |

Here we demonstrate that the HER model, in addition to autonomously learning hierarchically structured cognitive tasks, is able to account for a range of effects in dlPFC, mPFC, and their interactions previously reported in the literature. These effects are derived from signals in the model that compute differences between observed and predicted outcomes ^7,25^ (in mPFC), and signals related to the maintenance, updating, and modulation of working memory representations ^24,31^ (in dlPFC). Notably, all signals in the model derive from the computation of predictions errors, and thus suggest a common neural currency of error and error representation in prefrontal cortex.

## Results

### Context, Working Memory, & Control

The role of dlPFC in working memory and representation of task structure remains an ongoing research concern. In the past two decades, numerous fMRI studies have investigated the structure and function of dlPFC under various hierarchical task and working memory demands. In Koechlin et al. (2003)^24^, the authors investigated the function of dlPFC in two tasks while manipulating the amount of information conveyed by task-relevant stimuli. In their Motor Condition, activity throughout dlPFC – from areas labeled PMd (dorsal premotor cortex) to rostral dlPFC –was observed to increase monotonically as the information content of a contextual cue increased (fig 2, middle column). An additional increase in activity was observed only in PMd when subjects were required to make two responses rather than a single response. In Simulation 1 (Figure 2), the HER model accounts for the general trend of increasing activity across dlPFC as the increasing strength of error prediction representations learned by the model – more information means more potential errors that must be accounted for. This account complements the Information Cascade model ^24^ based on information theoretic formulations; in information theory, information is the amount by which uncertainty about a random variable decreases given another variable. Error predictions learned by the HER model are used to modulate outcome predictions in order to support correct behavior - that is, their role is to reduce uncertainty regarding the likely outcomes of actions. The HER model accounts for the additional increase in activity observed in PMd through the transient update of representations (see supplementary material) at the lowest model level when successive stimuli mandate different responses, while conditions in which only a single response is required do not entail an additional update (Fig. 2, left column, bottom).

**Figure 2.**
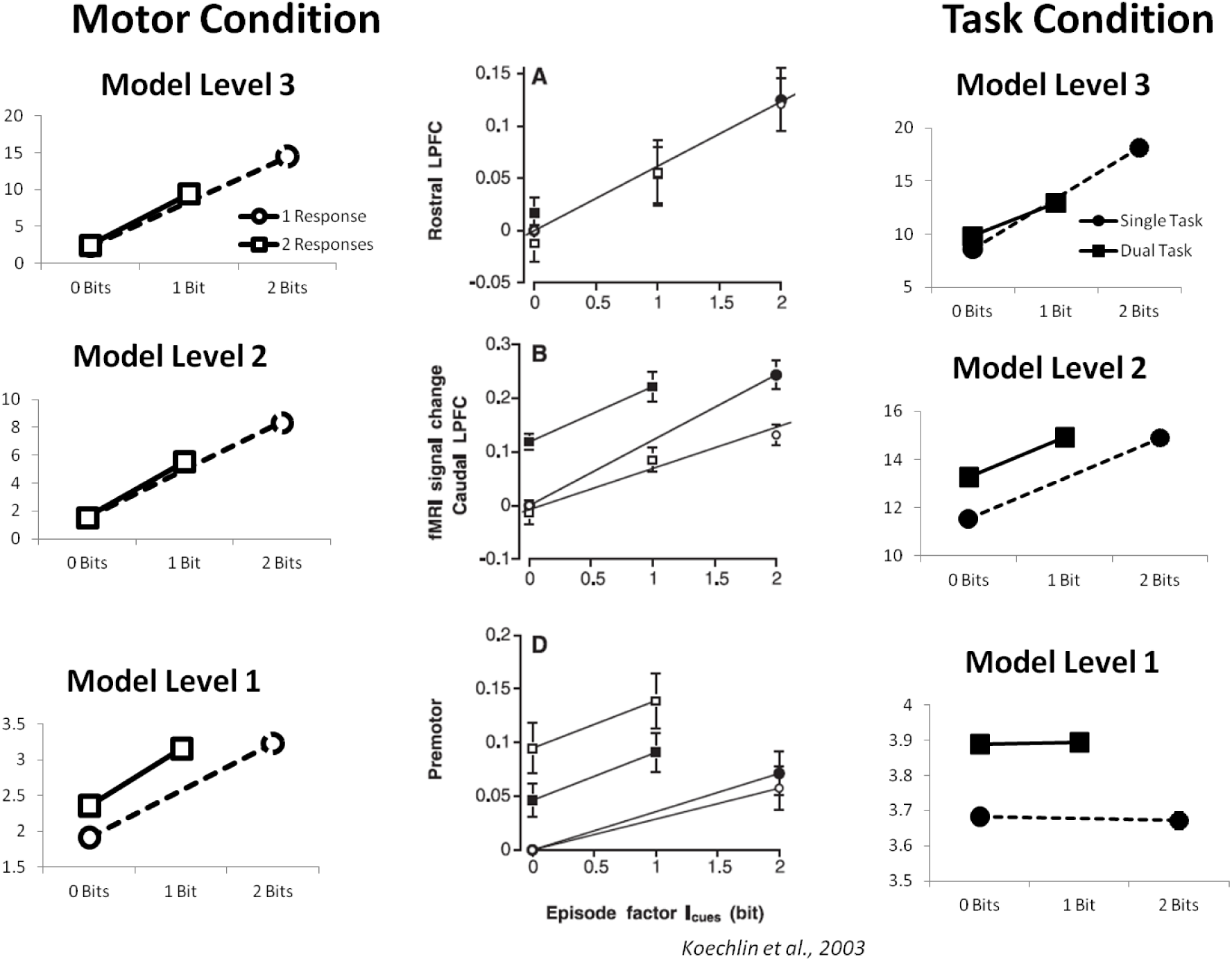
Information encoding in dlPFC. As the information content of a context cue increases, activity across hierarchically organized regions of dlPFC increases. The strength of error predictions maintained in dlPFC is proportional to information content: the more informative a cue is, the larger a reported error will be without the information supplied by that cue. The HER model captures effects of information related both to the nature of task-relevant stimuli (x axes) as well as responses that may be required (y axes). The HER thus provides a complementary account to the Information Cascade model of PFC. In the Task Condition of Koechlin et al.^24^ (right column), activity across dlPFC is observed to increase with the information content of a contextual cue. However, here activity in caudal dlPFC (middle frame) shows an additional increase when subjects must occasionally switch between two tasks (vowel/consonant, upper/lower case identification). This additional increase related to task switching is accounted for as transient increases in activity in the HER model when the nature of the task changes (Fig 2., right column, middle row).

### Learned Representation

While the HER model is able to capture a range of results related to the activity of ensembles of neurons reflected by the BOLD signal (supplementary material), it also posits a particular representation scheme deployed in dlPFC. Namely, single units in the HER model dlPFC each code for a component of a multi-dimensional error prediction. In addition to capturing data related to the strength of activity observed in dlPFC, then, the HER model should also be able to account for data relating to the activity of individual neurons as well as techniques designed to decode neural activity such as MVPA.

To investigate whether the error prediction representations learned by the HER model are consistent with those observed in human subjects, we recorded activity from the model as it performed the 1-2AX continuous performance task (Simulation 2, Figure 3A). We subsequently classified active representations in the model during periods of the task in which the model had been shown high- and low-level context variables (see Online Methods), but prior to a potential target cue being displayed. This approach is similar to the multi-voxel pattern analyses reported by Nee & Brown ^11^. Classification of the model representations is consistent with that observed in human subjects (Fig. 3A): at the lowest hierarchical level, sequences that may culminate in a target response (1A/2B) and those that will certainly not culminate in a target response (1B/2A) are represented in a distinct fashion (Fig. 3A, Bottom). However, the representations also partially overlap such that 1A sequences are partially categorized as 2B sequences, while 1B sequences are partially categorized as 2A sequences. At level 2 of the HER model, classification of each sequence is more decisive, with each unique sequence (1A/1B/2A/2B) being unambiguously decoded (Fig. 3A, Middle). This result is similar to human data, in which a region in mid-dlPFC shows a trend toward increased evidence for unique sequence coding. Finally, at the third hierarchical level (Fig. 3A, Top), sequences beginning with 1 or 2 are each collapsed (i.e.., equal evidence for 1A and 1B), reflecting the role of rostral dlPFC in coding high level context variables. The HER model explains the confusion of one target sequence with another (1A/2B) and one non-target sequence with another (1B/2A) at the lowest hierarchical level as a consequence of the increased activation of a predicted response common to both types of sequences – a target response in the former condition, and a non-target response in the latter condition.

**Figure 3.**
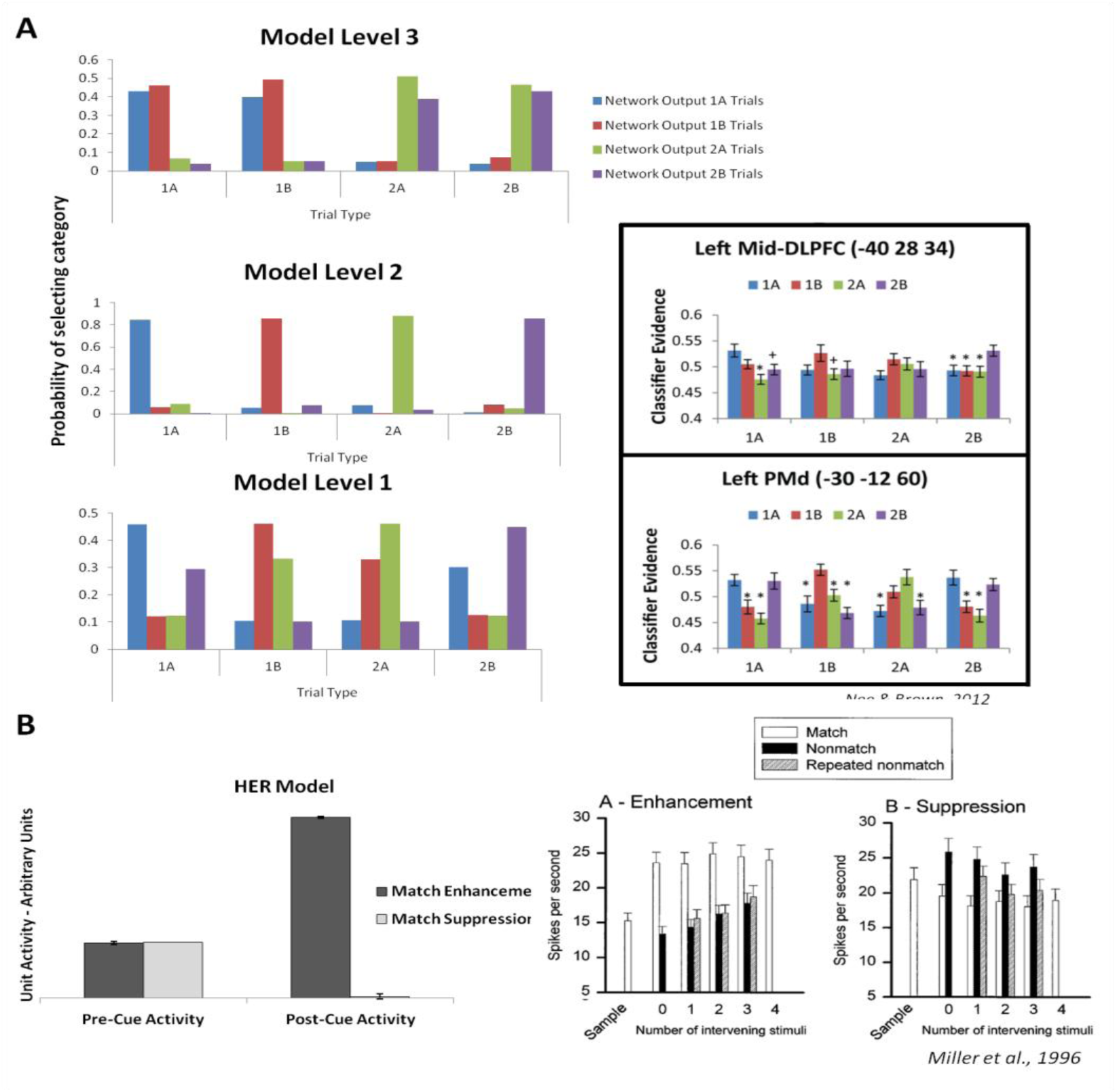
Distributed Representations in PFC. Separate units in the HER model represent components of a hierarchically-elaborated, multi-dimensional error prediction, suggesting how cognitive tasks may be represented neurally. A) Left: MVPA on error prediction representations maintained by the model while performing the 1-2AX CPT are consistent with human data showing that caudal regions of lPFC code for potential target sequences regardless of higher-order context, while more rostral regions encode more abstract context variables. Right: Human MVPA results, reprinted from Nee & Brown 2012^11^. Classification results of model representations are naturally more robust than pattern analysis of fMRI data since it is possible to record the activation of units in the model with perfect fidelity, while BOLD signals are subject to noise. B) Units in level 1 of the HER model (left) show activity related to match suppression and enhancement while performing a delayed match-to-sample task. Prior to observing a target stimulus, activity in these units reflects the equal probability of observing a match or non-match cue. Following the presentation of the target stimulus, the activity of units predicting the occurrence of a match is enhanced, while the activity for non-match-predicting units is suppressed, similar to data recorded from monkey lPFC (right). The HER model further predicts the existence of units showing effects of mismatch enhancement and suppression. Reprinted from Miller et al., 1996.

### Single-Unit Neurophysiology

The representation scheme proposed by the HER model suggests that individual neurons in lPFC should code for components of a distributed error representation, with single units signaling the identity and likelihood of observing a particular error. The model further suggests that these signals should evolve through the course of a trial as the likelihood of observing specific types of errors increases or decreases. We recorded activity in the model as it performed a delayed match-to-sample (DMTS) task (Simulation 3). Consistent with observed unit types recorded in macaque monkeys ^32^, units in the HER model were identified with increased activity following the occurrence of a target probe that matched the sample (match enhancement; Fig. 3B), while distinct units were identified whose activity decreased following a matching target (match suppression; Fig 3B). The HER model accounts for these two types of neurons as the modulation of predictions regarding possible responses following the presentation of a target cue. When a matching target is presented, the activity of units predicting a “match” response increases (enhancement) while the activity of units predicting a “non-match” response decreases (suppression). The HER model further suggests *a priori* that additional types of neurons should be observed in lPFC, namely *mismatch* enhancement and suppression neurons – neurons whose activity reflects the increased and decreased likelihood of making a non-match and match response, respectively.

### The neural bases of behavior in prefrontal cortex

In addition to reproducing effects from human fMRI data regarding the nature of stimulus representations in PFC, the HER model also suggests how these representations may influence patterns of behavior. In order to investigate the influence of hierarchically-organized representations on the timecourse of learned behaviors, we simulated the model (Simulation 4, Figure 4B) on a ternary probability estimation task ^33^ in which subjects were asked to estimate the probability that a compound stimulus, varying along two feature dimensions, belonged to each of three categories. Subjects were found to adopt three different strategies in their responses (Fig. 4B, bottom row): one group (Least Certain, LC, left) consistently assigned near-equal probabilities for each category, a second group (Label Margin, LM, center) assigned a low probability to one category and approximately equal probabilities to the other two, while the final group (Most Certain, MC, right) assigned a high probability to one category and low probabilities to the others. Similar patterns of behavior were observed in the HER model during simulated experiments in which the learning rate was manipulated as follows (Fig. 4B, top row). For simulations in which all learning was disabled, the model’s probability estimates corresponded to the LC group. When learning was enabled only for the lowest hierarchical level, the model’s behavior corresponds to the LM group, reflecting learned representations that allow the model to rule out one of the three categories but lacking the higher order information required to distinguish between the remaining two. Finally, when learning is enabled for all levels, the model rapidly learns the entire task, corresponding to the behavior of the MC group. In the HER model, these behaviors are intimately linked to learned error predictions: the model decomposes a task by selecting at each hierarchical level the stimulus feature that best reduces response uncertainty. The HER model thus provides an account of how neural representations acquired during learning might contribute to patterns of behavior.

**Figure 4.**
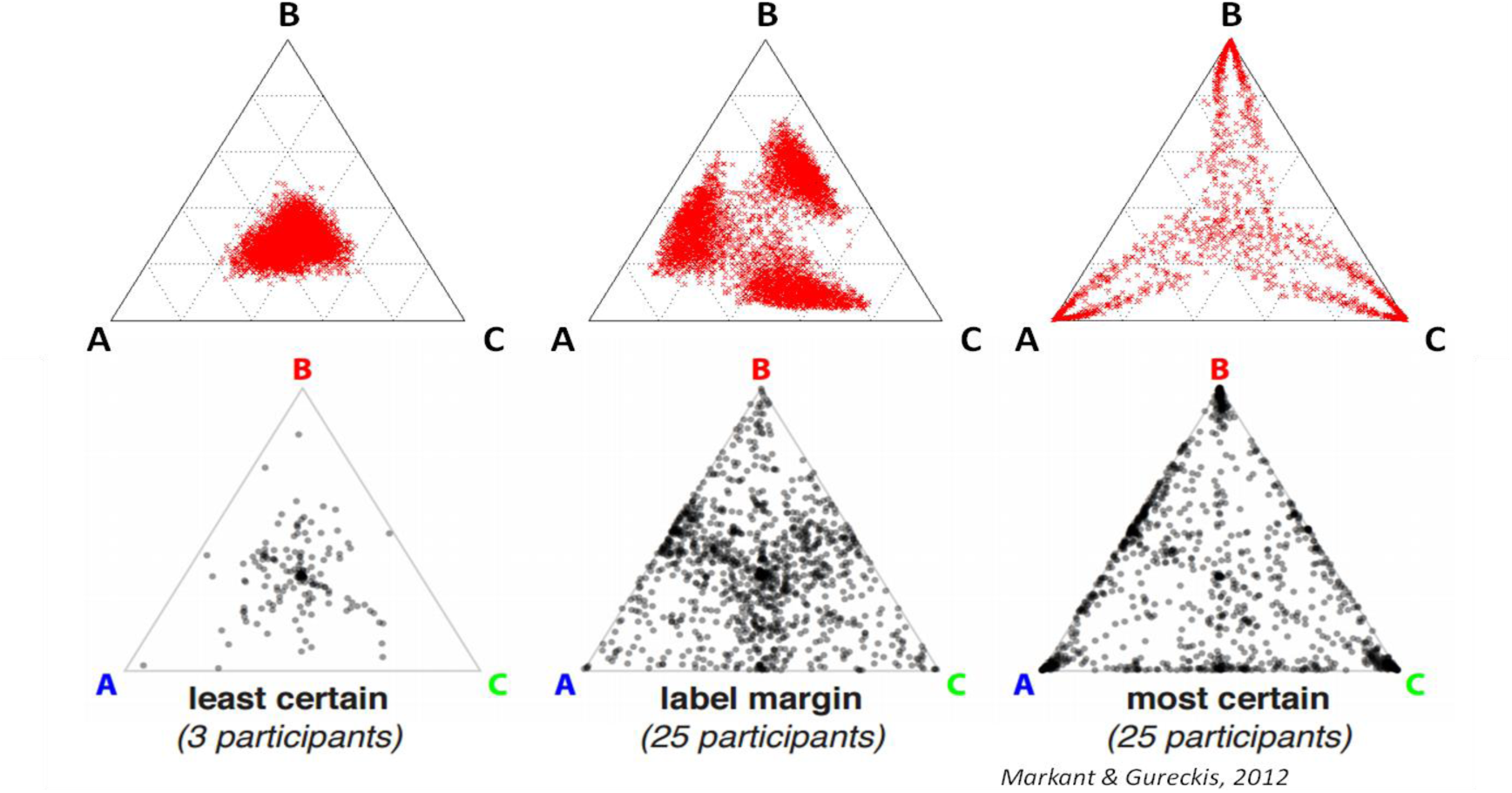
Connecting representations to behavior. Behavior of the HER model (top) with learning selectively enabled at zero (left), one (center), and all (right) hierarchical levels. The model’s estimate of the probabilities of three possible categories matches the behavior of three groups of human subjects (bottom) during a ternary probability estimation task. The HER model thus provides an account of how task representation at the level of single units contributes to behavior. Reprinted from Markant & Gureckis (2012)

### Interaction of mPFC and dlPFC

The HER model, being an extension of the PRO model of ACC/mPFC, already captures a wide array of effects observed within ACC^7,25^. The HER model extends the PRO model in two critical ways: first, it specifies how mPFC and dlPFC may interact in order to support sophisticated behaviors, and two, it suggests a parallel hierarchical organization of mPFC in which successive hierarchical regions report increasingly abstract error signals. Such an organization of mPFC has been proposed previously^34,35^, and, indeed, evidence has been found that supports a role for mPFC in processing hierarchical errors ^27^. The HER model is able to capture the pattern of activity observed by Kim et al. ^26^ (Simulation 5) for distinct regions of both mPFC and dlPFC (Fig. 5A, middle column). The HER model interprets activity in hierarchically-organized regions of mPFC as the discrepancy between increasingly abstract predicted and observed outcomes, consistent with the role of mPFC in error computation proposed by the PRO model ^7,25^, and complementary to the interpretation of Kim et al. However, while their notion of higher-order error signals is specified qualitatively, successively more abstract errors in the HER model are a product of quantitative predictions at lower levels that are insufficient to explain a subject’s observations, in line with the predictive coding framework that informs the structure of the HER model.

Additional evidence regarding the interaction of mPFC and dlPFC comes from studies of patients with dlPFC lesions ^36^. In a delayed match to sample task, an Error Related Negativity (ERN) is observed in subjects with lesions to dlPFC for both correct and incorrect trials (Fig. 5B, left column). The HER model (Simulation 6) explains this as the inability to maintain relevant information across a delay period in order to modulate predictions regarding likely outcomes (Fig 5B, right column). Without this additional contextual information available in the model, both correct and incorrect outcomes are surprising, resulting in increased mPFC activity in a lesioned version of the HER model on both types of trials.

**Figure 5.**
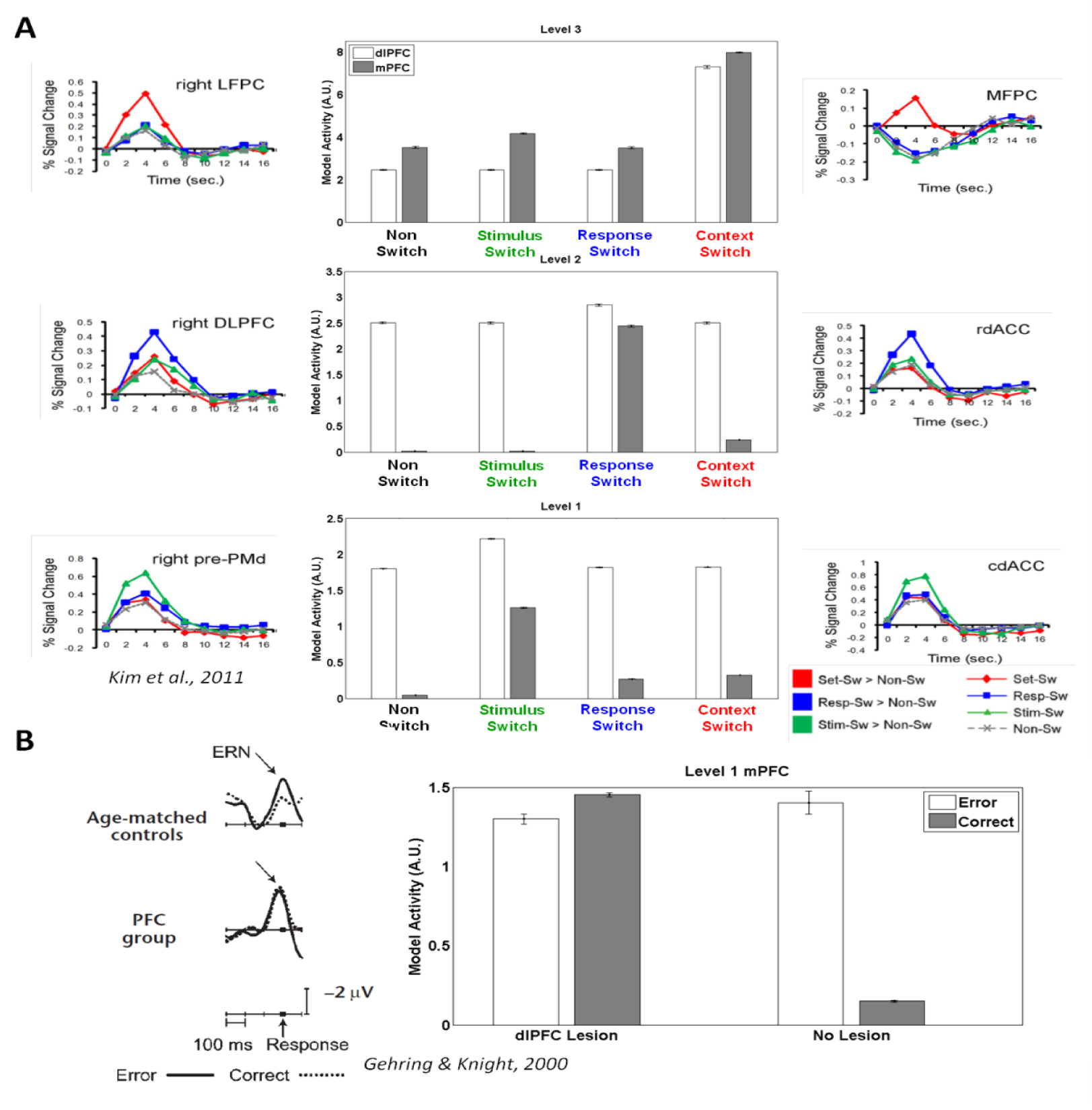
Interactions of mPFC and dlPFC. The HER model suggests how mPFC and dlPFC may cooperate to minimize prediction error through passing error and error prediction information through hierarchical levels. A) Increased activity in parallel hierarchical regions in the HER model, associated with mPFC and dlPFC, is associated with errors (mPFC) and updates of error predictions (dlPFC) at different levels of abstraction, from concrete (level 1, stimulus switch) to abstract (level 2, response switch; level 3, context switch). B) Modulation of mPFC by error predictions maintained in dlPFC is critical for contextualizing predictions regarding the likely outcome of actions. In a delayed-match-to-sample task, the HER model correctly captures the elimination of the ERN following correct trials due to the maintenance of information regarding the sample cue. However, when the model is lesioned such that information normally maintained in dlPFC is no longer available to mPFC, the model produces an ERN to correct and error trials alike.

### Application to Machine Learning

Recent advances have led to machine learning algorithms, such as deep learning, capable of performing complex tasks at or above human level. While the HER model has previously been shown to learn tasks used to investigate human cognition at a level comparable to deep learning algorithms ^28^, a stronger test involves examining the model’s performance on typical machine learning tasks. We therefore simulated a slightly modified version of the HER model (Simulation 7, See Online Methods) on a suite of tasks previously used as benchmarks for investigating the performance of machine learning algorithms^37,38^. The results of these simulations (Table 1) are notable for two reasons. First, while no particular effort was made to optimize model parameters, the HER framework compares favorably to several out-of-the-box ML algorithms on a number of benchmark tasks. Additionally, the HER model performs these tasks using the same general architecture as used in our simulations of human behavior and neural activity, indicating that the HER model provides a general framework that may scale well and be applied to a wide range of problems with little or no modification.

### Additional simulations

The results reported above are by no means exhaustive, but rather were selected to highlight how the HER model is able to account for patterns of activity observed in dlPFC and mPFC. Simulations of additional tasks are included in supplementary online materials. The results of these additional simulations show how the HER model simulates a variety of other fMRI findings. These serve to emphasize the main point that the HER model of PFC, as an instance of predictive coding formulations, is able to autonomously learn complex tasks in a manner that reproduces patterns of behavior, neuropsychological effects, and neural activity as measured by fMRI, EEG, single unit neurophysiology observed in empirical investigation.

## Discussion

In this paper, we have deployed a new computational neural model, consistent with known anatomy ^29,30^, to simulate a range of effects observed in studies of mPFC and dlPFC. Simulations demonstrate that the HER model captures various dlPFC effects, as well as how dlPFC and mPFC interact to support the acquisition and execution of sophisticated cognitive tasks. Because the HER model extends our previous PRO model of ACC/mPFC ^7^, it can also comprehensively account for mPFC activity in simple cognitive control experiments as previously reported ^7,25^. These results, taken as a whole, make the HER model among the most comprehensive models of PFC to date and provide an existence proof that simple predictive coding can account for a large corpus of PFC empirical findings.

The HER model provides a complementary perspective on existing models. Donoso et al. ^39^ cast the PFC as searching for, evaluating, selecting, and discarding task strategies to maximize reward. In the HER model, task strategies are represented automatically as hierarchical self-organized abstract representations of task context, which serve as a working memory basis for guiding behavior. Strategies are discarded from working memory when they no longer provide useful predictive information about subsequent events, or when contingencies change such that predictive information in working memory is repurposed by retraining its connections to modulate lower level predictions differently. The HER model can switch strategies flexibly as task cues change, and it can learn new responses when environmental contingencies changes. As with other neural models that include PFC ^40^, as well as models of hierarchical behavior ^4,41^, the HER model captures key aspects of neural anatomy, neurophysiology, and behavior during performance of cognitive tasks. The HER model further addresses the question of how these tasks might be learned in the first place, as well as how the components of a task are represented as expected prediction errors. The HER model thus fills a critical void left by models concerned with how coherent behaviors are organized based on pre-existing representations without specifying the nature of those representations or how those representations were acquired ^4,40,41^.

More generally, the HER model demonstrates how the predictive coding framework may be extended into prefrontal cortex in order to account for sophisticated cognitive behaviors. Each hierarchical level of the HER model is a relatively straightforward RL learner based on previous models of mPFC ^7,25^, and augmented with a WM component able to maintain representations over periods of time. It is notable that the model is not only able to replicate effects observed throughout PFC during the performance of complex tasks, it learns these tasks autonomously in a manner comparable to human performance^28^, despite its simple motif structure.

## Acknowledgments

WHA and JWB developed the HER model. WHA implemented the model, conducted simulations and analysed data. WHA and JWB wrote the manuscript. WHA was supported in part by FWO-Flanders Odysseus II Award #G.OC44.13N. The authors wish to thank Tom Verguts, Clay Holroyd, Eliana Vassena, Matthew Botvinick, Maynard James Keenan, Derek Nee and Todd Braver for helpful comments and discussion in the preparation of this manuscript. Supported in part by the Intelligence Advanced Research Projects Activity (IARPA) via Department of the Interior (DOI) contract number D10PC20023. The U.S. Government is authorized to reproduce and distribute reprints for Governmental purposes notwithstanding any copyright annotation thereon. The views and conclusions contained herein are those of the authors and should not be interpreted as necessarily representing the official policies or endorsements, either expressed or implied, of IARPA, DOI or the U.S. Government.

## Methods

### The Hierarchical Error Representation Model

A detailed description of the HER model is provided in our previous publication^1^. Here we provide a summary of the components of the HER model and how they interact. The HER model is composed of multiple levels, each instantiating a relatively simple RL learner based on the PRO model of mPFC^2^, and endowed with a working memory gating mechanism that governs whether a stimulus is stored in WM or not. Levels interact with one another through top-down and bottom-up pathways.

### Reinforcement Learning

The output of each level is determined by the item currently stored in WM at each level and the strength of weights associated with that item:

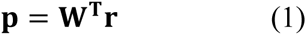

where **p** is a vector of predicted outcomes, **r** is the item currently stored in WM, and **W** is a weight matrix associating **r** and **p**. Errors at each level are computed as the difference between observed and predicted outcomes:

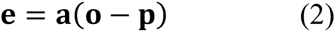

where **o** is the observed outcome and **a** is a filter set to 0 for unselected actions and 1 everywhere else, effectively preventing learning about unselected actions in the model. Weights are updated according to:

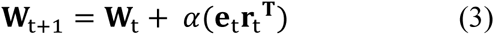

where is a learning rate parameter and *t* indicates the current model iteration.

### Working Memory Gating

The WM gating mechanism, inspired by models of basal ganglia^3^ determines whether a currently presented stimulus will be stored in WM at each level. This determination is made based on the learned, relative value of encoding a new stimulus in WM vs. maintaining the current contents of WM:

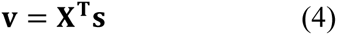

Here **s** indicates a vector of external stimulus features (distinct from internal representations), **X** is a weight matrix associating stimulus features with WM representations, and **v** is the value of storing a particular feature in **s** as a representation in working memory **r**. Weights **X** are trained through backpropagation of the error term calculated in eq. 2:

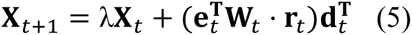

While error backpropagation of error is biologically implausible, our intent was not to develop a neurally faithful model of basal ganglia, and so backpropagation was selected for computational convenience. Nevertheless, in our previous work, we demonstrate how a more realistic model of WM gating using scalar reinforcement signals may be implemented in a manner consistent with previous proposals ^3–5^, so that the HER model functions as well while maintaining biological plausibility. The value of storing stimulus features in WM is passed through a softmax function in order to determine whether a stimulus will be stored in WM or the current contents of WM will be maintained.

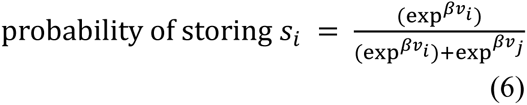

is a gain parameter governing the probability of selecting the highest value in **v**.

### Bottom-Up and Top-Down Pathways

The RL algorithm and WM gating mechanisms operate at each level of the HER model. Layers interact with one another through bottom-up and top-down pathways based on Predictive Coding formulations. In the bottom-up pathway, errors reported by a given hierarchical level (eq. 2) are passed to a superior hierarchical level. In the HER model, the error reported by a level, conjoined with active WM representations at that level, acts as the outcome for the next higher level:

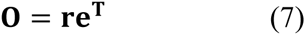

Here **O** is the matrix computed from the outer product of the error and WM representation vectors. For computational convenience, this is reshaped into a vector **o**. Error computation and learning is identical at each level (eqs 2 & 3), with the exception that the outcome term at each level above the 1st is derived from eq. 7.

The purpose of training higher-order levels in the hierarchy using the outcome term in eq. 7 is to derive predictions regarding the likely errors that can be expected at lower-order levels. Predictions at each level are calculated as in eq. 1; however, at higher-order levels, these predictions reflect expected errors reported by lower levels. Since knowledge of likely errors can be useful in avoiding those errors, the predictions generated at each level can be used to modulate the predictions generated by inferior levels. For levels above the 1st, the prediction **p** is reshaped into a matrix **P** of the same dimensionality as the weight matrix **W** of the immediately inferior level. **P** and **W** are then added to one another, resulting in a modulated weight matrix used to compute a prediction of likely outcomes that incorporates higher-order information:

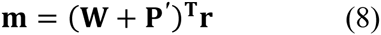

where **m** is the modulated prediction.

### Responses

At the base level, model activity is translated into response probabilities. As in the PRO model^2^, the HER model learns predictions of response-outcome associations. Individual responses may be associated with either correct or error feedback. In order to generate a response, the learned likelihood of receiving correct feedback is compared to the learned likelihood of receiving error feedback for each candidate response:

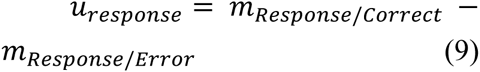

which is then passed through a softmax function to determine the probability of the model making each response:

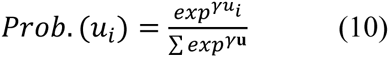

where is a gain parameter.

### Sources of Activity in Prefrontal Cortex

A key concern in relating computational neural models to empirical data, especially with regard to indirect measures of neural activity such as EEG and BOLD signals, is in selecting an appropriate measure of model activity. In the PRO model^6^, ACC/mPFC activity was interpreted as negative surprise – the ongoing difference of predicted outcomes minus actual observations. In the HER model, the role of mPFC is identical to its function in the PRO model, and thus the measure of model activity for mPFC remains the same:

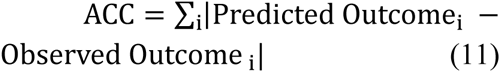

In comparison, dlPFC is thought to have multiple underlying mechanisms that contribute to its temporal activity profile. First, and central to its role in working memory, sustained dlPFC activity is observed during maintenance periods of a task when one or more items must be remembered in order to inform future behavioral responses. Second, the process of encoding an item in WM corresponds to a transient increase in bold activity following the presentation of an item to be maintained in WM. Finally, the level of sustained activity observed in dlPFC is additionally modulated by higher order information.

The role of dlPFC in the HER model is to learn to represent task stimuli that reliably precede prediction error signals generated by ACC. That is, dlPFC learns the expected error given a stimulus S:

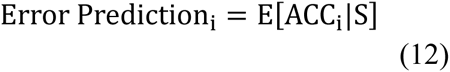

Using Error Prediction as the unit of currency, then, we model the three sources of dlPFC activity described above as follows. First, sustained dlPFC activity related to WM maintenance is calculated as the absolute value of active error predictions:

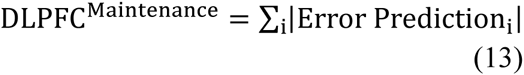

Second, transient activity related to updating the contents of WM is modeled as the absolute difference on successive model iterations, t-1 and t, of active error predictions:

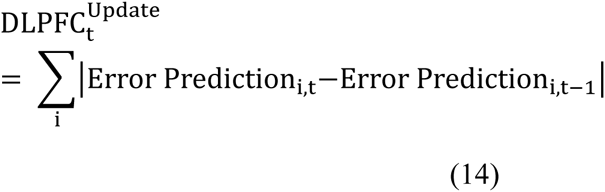

Finally, the influence of top-down information on sustained activity is modeled as the difference between the Error Prediction for a given level, and the Error Prediction at that level if there were no top-down modulation of error predictions:

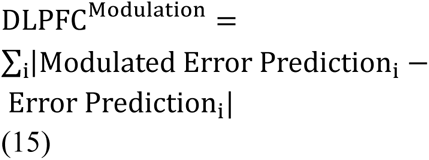

While it is likely that these three sources contribute to dlPFC activity in differing measures, we remain agnostic as to their relative contributions. Therefore, in order to compute an overall measure of dlPFC activity in the model, eqs 13-15 are simply summed together:

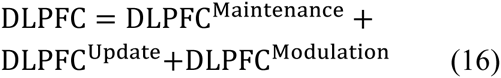

Using eq. 16 as our measure of dlPFC activity, we show that the HER model is able to reproduce patterns of activity observed in hierarchically organized regions of dlPFC for tasks involving significant WM demand. Moreover, we show that the proposed role of dlPFC in learning and maintaining representations of error is sufficient to reproduce data from single-unit and MVPA studies investigating the nature of representations in dlPFC. Finally, using eq. 11 as our measure of mPFC activity, the HER model is able to capture the joint pattern of activity observed in mPFC and dlPFC data.

## Simulations

In order to ensure that the effects reported were not due to a specific parameterization tailored to each task, all simulations were conducted using a common parameter set (table 1) unless otherwise noted below. In previous work^1^ we examined the influence of alternative parameterizations on the development of the HER model. Parameters for the current simulations were chosen based on this prior exploration in order to ensure that the model was able to learn each task.

### Simulation 1: Koechlin et al (2003).

We simulated the HER model on the behavioral task described in Koechlin (2003)^7^. For the sake of brevity, we refer the reader to the supplementary online material associated with that paper for a detailed description. Briefly, subjects participated in two experiments, a “motor” experiment and a “task” experiment. In each experiment, the subjects experienced blocks of 12 sequentially presented stimuli whose visual appearance(color) indicated which response they were to make (in the case of the motor experiment) or which task (vowel/consonant or upper/lower case discrimination) they were to perform. Each block was preceded by a context cue which indicated the mapping between stimulus color and the appropriate response or task. We simulated the motor and task conditions separately. For the “motor” experiment, inputs to the model were the 4 context cues associated with each condition, and 6 colors that were observed by the subjects during the experiment. 2 responses were possible (left or right), and feedback to the model indicated either correct or incorrect performance. For the “task” experiment, model inputs were the 4 context cues associated with each condition, 6 colors observed during the experiment, and 4 cues indicating whether the stimulus was a vowel or consonant, or upper or lower case. 4 responses could be generated by the model, indicating upper or lower case or vowel or consonant responses. Feedback to the model indicated correct or incorrect performance. The model performed each condition for 4500 blocks (1 block = 12 trials).

### Simulation 2: 1-2AX CPT

The 1-2AX CPT^3,8^ is a hierarchically organized task in which a subject’s response to a target cue (‘X’ or ‘Y’) is governed by both the cue that immediately preceded it (’A’ or ‘B’), as well as a “context” cue (’1’ or ‘2’) that indicates which target sequence (’AX’ or ‘BY’) is valid at any given time. Sequences of stimuli may be thought of as being organized in ‘inner’ and ‘outer’ loops, where inner loops are composed of 2-stimulus sequences with ’A’ or ’B’ followed by ’X’ or ’Y’, and outer loops are the sequence of inner loops followed by the presentation of a context cue. We simulated the HER model on a version of the 1-2AX task as described in O’Reilly & Frank (2006)^3^ in which each outer loop consisted of 1-4 inner loops, and the probability of observing a valid sequence was 0.25. There were 8 inputs to the model, corresponding to the 6 relevant cues in the task, as well as 2 distractor cues that had no task relevance. At each cue, the model made a response to indicate whether the current stimulus was a target or not. In order to perform the task correctly, target responses should be made only at the presentation of a valid target cue; all other cues should result in non-target responses. Feedback to the model indicated correct or incorrect performance. We simulated the 1-2AX task on approximately 24,000 individual cue presentations as described in previous work^1^. The activity of each prediction unit at each higher level was recorded on the presentation of a potential target cue (’X’ or ’Y’) to be used as input to a 2 level feedforward neural network with 10 hidden units. The neural network was trained on the sequence of high (’1’ & ’2’) and low (’A’ & ’B’) level context cues using the MATLAB ^9^ neural networks toolbox.

### Simulation 3: Miller, Erickson, and Desimone (1996)

The model was simulated for 6000 trials on a simple delayed match-to-sample (DMTS) task. On each trial, a neutral stimulus indicating the beginning of a trial was presented, followed by one of two sample stimuli, and ending with one of two target stimuli. There were a total of 3 inputs to the model, 1 indicating the trial onset, and 2 for the task-relevant stimuli. Note that sample stimuli and target stimuli used the same representation. The model could make two responses indicating either a match or non-match between the sample and target stimuli, and each response resulted in either correct or incorrect feedback, for a total of 4 outcome units.

### Simulation 4: Markant & Gureckis (2012)

The model was simulated on a ternary probability estimation task for 5000 trials in three different learning conditions. Task stimuli were modeled as compound stimuli composed of two feature dimensions, and each dimension had three possible values as described in previous work^1^. Each unique conjunction of feature dimension values was associated with one of three possible responses such that each feature of each dimension was associated with each of the three possible responses in only one instance. The learning rate of the model was manipulated across the three learning conditions as follows: in the no learning condition, the learning rate for all hierarchical levels of the model was set to 0, and thus no learning occurred during the experiment. In a second learning condition, learning was enabled only for the lowest hierarchical level and set to the parameter values reported in table 1. Finally, in the third learning condition, learning was enabled for all levels, and set to the parameter values in table 1.

### Simulation 5: Kim et al. (2011)

In Kim et al. (2011)^10^, the authors attempted to identify brain activity related to set switches at various levels of abstraction. Subjects were presented with a colored (red or green) box situated in a single cell of a 2X2 grid. The color of the box indicated a cognitive set of two numbers that the box might indicate; a red box, for instance, may indicate either 5 or 7, depending on the box’s horizontal position (left/right columns). A red box appearing in the left column may indicate a 5, while a red box in the right column indicates a 7. The vertical position of the box (upper/lower rows) indicated which of two operations (greater than/less than) the subject should perform in comparing the number indicated by the position and color of the box with a plain digit presented alongside the box. Changes in the position and color of the box indicate set switches at various levels of abstraction: changes in the left/right position indicate stimulus switches, changes in the upper/lower position indicate switches in responses, and changes in box color indicate cognitive set switches. The model was simulated on 20,000 trials, and activity was calculated from the final 2000 trials. A total of 6 inputs were modeled, reflecting the 2 colors, 2 horizontal positions, and 2 vertical positions possible in the task.

### Simulation 6: Gehring & Knight (2000)

The model was simulated for 6000 trials on a simple delayed match-to-sample (DMTS) task. On each trial, a neutral stimulus indicating the beginning of a trial was presented, followed by one of two sample stimuli, and ending with one of two target stimuli. There were a total of 3 inputs to the model, 1 indicating the trial onset, and 2 for the task-relevant stimuli. Note that sample stimuli and target stimuli used the same representation. The model could make two responses indicating either a match or nonmatch between the sample and target stimuli, and each response resulted in either correct or incorrect feedback, for a total of 4 outcome units. Two conditions were simulated: a control condition in which all pathways were intact, and a lesion condition in which the value of P’ in Eq. 8 was set to 0, effectively removing any top-down influence between levels. Activity was recorded from all 6000 trials.

### Simulation 7: Machine Learning

In order to assess the suitability of the HER model for typical machine learning applications, the model was modified as follows. First, the WM gating mechanism was bypassed, and the contents of WM at each level were set to the input the array of features appropriate to each data set. This deviates from the HER model insofar as, under the machine learning simulations, multiple values (instead of a single value) could be stored in WM on a single trial. Additionally, the values of WM representations were continuous numbers rather than binary as in our simulations of neural data. The output of the model was passed through a sigmoid activation function rather than using a softmax function to determine the models output. The model was simulated on a range of tasks reported in Meyer et al. (2003)^11^ using the data sets described in that paper and consisting of 100 training and test sets under a 10 times repeated 10-fold cross validation regime. For each task, the model was trained on each training set for 100 epochs (1 epoch = 1 pass through the training data), and validation on the test set was performed after every second epoch. Model performance assessed using the percent error calculated in two ways: first as the minimum of the mean/median of the error for 100 epochs (1 pass through training data) for all 100 training sets. Error was also calculated as the mean/median of the minimum error over 100 training epochs for all 100 training sets. This second approach treats the test data for each training set as a validation set, and thus limits how well the network may generalize to novel data.

**Table 1:**
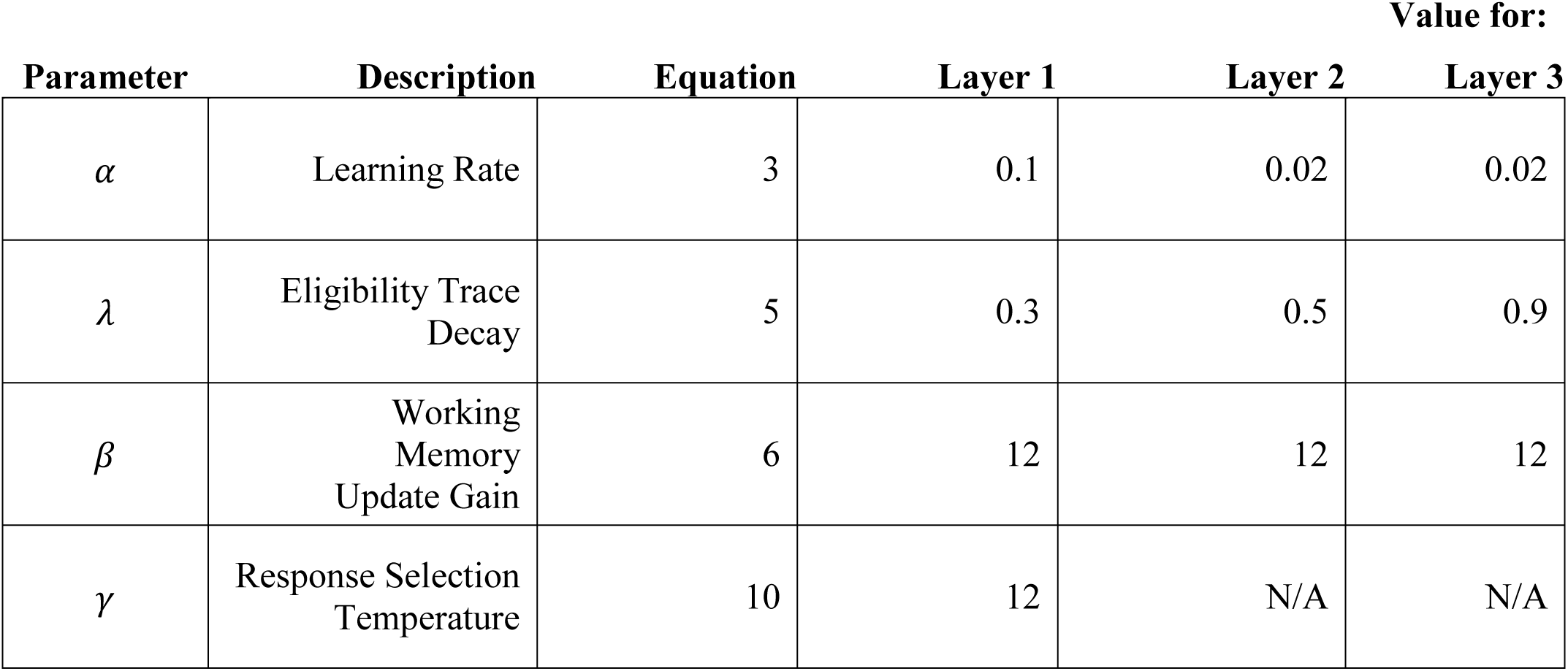
Parameter set for all simulations

